# Mutations within the Open Reading Frame (ORF) including Ochre stop codon of the Surface Glycoprotein gene of SARS-CoV-2 virus erase potential seed location motifs of human non-coding microRNAs

**DOI:** 10.1101/2021.06.19.449095

**Authors:** Krishna Himmatbhai Goyani, Shalin Vaniawala, Pratap Narayan Mukhopadhyaya

## Abstract

MicroRNA are short and non-coding RNA, 18-25 nucleotides in length. They are produced at the early stage of viral infection. The roles played by cellular miRNAs and miRNA-mediated gene-silencing in the COVID-19 epidemic period is critical in order to develop novel therapeutics. We analysed SARS-CoV-2 Surface Glycoprotein (S) nucleotide sequence originating from India as well as Iran, Australia, Germany, Italy, Russia, China, Japan and Turkey and identified mutation in potential seed location of several human miRNA. Seventy single nucleotide polymorphisms (SNP) were detected in the S gene out of which, 36, 32 and 2 were cases of transitions, transversions and deletions respectively. Eleven human miRNA targets were identified on the reference S gene sequence with a score >80 in the miRDB database. Mutation A845S erased a common binding site of 7 human miRNA (miR-195-5p, miR-16-5p, miR-15b-5p, miR-15a-5p, miR-497-5p, miR-424-5p and miR-6838-5p). A synonymous mutation altered the wild type Ochre stop codon within the S gene sequence (Italy) to Opal thereby changing the seed sequence of miR-511-3p. Similar (synonymous) mutations were detected at amino acid position 659 and 1116 of the S gene where amino acids serine and threonine were retained, abolishing potential seed location for miR-219a-1-3p and miR-20b-3p respectively. The significance of this finding in reference to the strategy to use synthetic miRNA combinations as a novel therapeutic tool is discussed.

## Introduction

The recent epidemic of the novel coronavirus disease 2019 or the COVID-19 is caused by a new coronavirus that was detected in December 2019 in the Wuhan city of China. Thereafter it has rapidly spread across the world like wildfire and turned into a global pandemic. [1] Soon after the pandemic emerged the causal pathogen was identified as a Coronavirus and later named SARS-CoV-2. [1]

The Genus Coronaviruses, abbreviated as CoVs, are classified into the family Coronaviridae and sub family Coronavirinae. These viruses are characterized by typical crown-like spikes on their surface (Latin: Corona = Crown) and hence the name ‘’Coronavirus’’. [2] These viruses infect humans, birds and other mammals and the infection causes a varied degree of complexities ranging from respiratory, neurological, enteric to hepatic diseases. [3]

The genetic material of Coronaviruses is an unsegmented single stranded ribonucleic acid (RNA) with a size that ranges from 26 to 32 kilobases in length. Serological and genomic differences have further divided the Coronavirinae subfamily into four major genera, namely Alphacoronavirus, Betacoronavirus, Gammacoronavirus, and Deltacoronavirus. [3] While the former primarily infect mammals, the latter two are known to infect birds. [4]

SARS-CoV-2 belongs to the Betacoronavirus family. This is the 7^th^ Coronavirus that has been recorded to infect humans. Among these seven, four of them (229E, NL63, OC43, and HKU1) cause mild symptoms of cough and cold. On the other hand, the other 3 Coronaviruses (SARS-CoV, MERS-CoV, and SARS-CoV-2) cause serious symptoms that are associated with mortality at the rate of 10%, 37%and 5% respectively. Since the discovery of SARS-CoV-2 a large amount of research work, studies and trials has been underway. [5,6] Yet, no viable and full-proof treatment protocol has emerged for SARS-CoV-2 infections that can improve the outcome in patients. [7]

The entire single stranded RNA genome [8] has been characterized by applying high-end RNA-based metagenomic, passive parallel sequencing technology and found to measure 29,881 bp in length (GenBank accession number MN908947) that encodes 9860 amino acids. [9] The genes present within this genome code for both structural as well as non-structural proteins. While structural proteins are coded by the S, E, M and N genes, the non-structural proteins of 3-chymotrypsin like protease, papain-like protease, and RNA-dependent RNA polymerase are expressed from within the ORF region of the genome. [10]

The SARS-CoV-2 viral particle is covered by a large number of glycosylated S proteins. These proteins attach to the host cell receptor angiotensin converting enzyme 2 (ACE2) thus facilitating the entry of the virus into the human system. [11] Once the S protein attaches to its receptor, the Transmembrane (TM) protease serine 2 or TMPRSS2 (a type 2 TM serine protease) that is positioned on the cell membrane of the host, assists the entry of the virus into the cell by activating the S protein. Upon entry into the cell, the virus releases the RNA that soon replicates resulting in synthesis of structural proteins which are then assembled and packaged to create and release more viral particles. [12]

S proteins are essential and crucial for the completion of the viral life cycle and hence provide potential targets for therapy. For example, recently, a novel vinylsulfone protease inhibitor has been shown to be effective against SARS-CoV-2. [13] An important feature of the SARS-CoV-2 S protein is its high level of conservation across all human Coronaviruses (HCoVs) and in all cases, its role lies in recognition of the receptor, assisting the virus in attaching with it and facilitating entry of the virus within the cell. Needless to say, the S protein therefore is an intense object of study and analysis in recent times.

MicroRNA or the miRNA are species of ribonucleic acid molecules that do not code for any protein. They function like regulators by attaching to the messenger RNA and influencing the translation process. A large number of research-based evidences has come up that demonstrate that human miRNA interacts with pathogenic viral RNA and adversely influence their pathogenesis. [14,15]

The mode of action of the cellular miRNA in acting against viral RNA is not fully understood. Nevertheless, it is well studied that they directly interfere with viral replication inside the cells. One proposed mechanism is inhibiting viral RNA translation by sudden deregulation of a vast number of cellular miRNAs that collectively bind and inhibit viral RNA processing. [16,17] This antiviral response in turn affects the cellular mRNA which are bonafide targets of these specific human miRNAs. Such mRNA are members of the cell signalling pathways that participate in the cellular response sparked off by the viral infection, such as WNT, INF, PIK3/AKT, MAPK, and NOTCH.[18] On the other hand, other process such as skipping of crucial binding of these cellular miRNA, error in synthesis from the parent DNA and such others can favour viral replication and subsequent expansion of the pathogen population inside the body and establishment of the disease. [19]

In this study, we analysed mutations within the nucleotide sequences of the S gene of SARS CoV-2 that were submitted in the public database and performed in-depth bioinformatic analysis using advanced miRNA analytical tools to predict potential alteration in binding of human miRNA to this gene. We hypothesize that over time, various transitions and transversions surfaced in the SARS-CoV-2 S gene that positively benefited the pathogen in evading host miRNA-mediated inactivation resulting in establishment of the disease. Our finding identifies a potential cause for varying degrees of COVID-19 disease progression in the population given that these mutations were not ancient, yet and unlike the D614G mutation for example, which is found in most of the isolated strains at the later stage of the pandemic and is known to reduce S1 shedding of the virus that increases infectivity. [20]

## Materials and Methods

The published nucleotide sequences of the SARS-CoV-2 ‘’S’’ gene coding for Spike glycoprotein were collected from the web resources of the National Center for Biotechnology Information – Nucleotide database.

The nucleotide sequence with accession number NC_045512.2 and titled ‘severe acute respiratory syndrome coronavirus 2 isolate Wuhan-Hu-1, complete genome’ was considered as the reference sequence for bioinformatic analysis. Downloaded nucleotide sequences were aligned with the reference using the Clustal Omega online tool on web servers accessed from the website ‘Mobyle@Pasteur’ or the ‘EBI web server’. [21] This is a multiple sequence alignment program that uses seeded guide trees and hidden Markov Model (HMM) profile techniques to generate alignments. Mutations were located based on variation observed in the test sequences when compared with the reference. S glycoprotein annotations were obtained from the website of the UniProt Consortium. [22] Human miRNA with potential seed sequence by which it can bind to the SARS-CoV-2 S RNA was detected using the miRDB tool. [23] Seed sequences with known mutations within the surface glycoprotein gene of SARS-CoV-2 were manually detected and listed.

## Results

A set of 31 nucleotide sequences for the surface glycoprotein gene of SARS CoV-2 virus submitted from India and with collection dates ranging from the period of 11-06-2020 to 01-04-2021 were analysed. It also included representative nucleotide sequences of this gene from the countries of Iran (n=2), Australia (n=5), Germany (n=5), Italy (n=5), Russia (n=4), USA (n=5), China(n=6), Japan (n=5) and Turkey (n=4). Seventy single nucleotide polymorphisms (SNP) were detected within all the S gene nucleotide sequences analysed. Out of them,36 involved transitions and were interchanges of two-ring purines (A to G) or of one-ring pyrimidines (C to T). Remaining 32 SNPs were transversions with interchanges of purine for pyrimidine bases, which involve exchange of one-ring and two-ring structures. Fourteen mutations were found to be synonymous (no change in amino acid) while 54 were found to be non-synonymous (change in amino acid). Two events of deletions were also detected. The details are summarized in Table 1.

**Table 1:** Details of SARS-CoV-2 Surface glycoprotein gene analysed in this study.

| Sr.no | GenebankAccession no. | Nucleotide position<sup>a</sup> | Nucleotide position<sup>b</sup> | Mutation<sup>c</sup> | Codon change | Mutation<sup>d</sup> | Laboratory mutation Code | Origin of sequence<sup>e</sup> | Release date<sup>f</sup> | Collection date<sup>g</sup> |
| --- | --- | --- | --- | --- | --- | --- | --- | --- | --- | --- |
| 1 | MZ268208.1 | 22394 | 882 | C>T | GAC>GAT | p.Asp294Asp | SM -26 | India/Gandhinagar | 24-05-2021 | 28-11-2020 |
|  |  | 23353 | 1841 | A>G | GAT>GGT | P.Asp614Gly | SM-39 |  |  |  |
|  |  | 23537 | 2025 | G>T | CAG>CAT | p.Gln675His | SM-42 |  |  |  |
|  |  | 23575 | 2063 | C>T | GCT>GTT | p.Ala688Val | SM-47 |  |  |  |
|  |  | 25302 | 3790 | G>T | GTG>TTG | p.Val1264Leu | SM-69 |  |  |  |
|  |  | 24876 | 3364 | G>T | GTG>TTG | p.Val1264Leu | SM-65 |  |  |  |
| 2 | MZ268209.1 | 21580 | 68 | A>G | CAA>CGA | p.Gln23Arg | SM-05 | India/Gandhinagar | 24-05-2021 | 28-11-2020 |
|  |  | 21924 | 412 | G>T | GAT>TAT | p.AsP138Tyr | SM-15 |  |  |  |
|  |  | 22886 | 1374 | G>T | AAG>AAT | p.lys458Ile | SM-33 |  |  |  |
|  |  | 22914 | 1402 | A>G | ATT>GTT | p.Ile468Val | SM-34 |  |  |  |
|  |  | 23353 | 1841 | A>G | GAT>GGT | p.Asp614Gly | SM-39 |  |  |  |
| 3 | MZ268599.1 | 21657 | 145 | C>T | CAT>TAT | p.His49Tyr | SM -08 | India/Gandhinagar | 24-05-2021 | 29-11-2020 |
|  |  | 22394 | 882 | C>T | GAC>GAT | p.Asp294Asp | SM -26 |  |  |  |
|  |  | 23353 | 1841 | A>G | GAT>GGT | p.Asp614Gly | SM-39 |  |  |  |
|  |  | 23543 | 2031 | G>T | CAG>CAT | p.Gln677His | SM-44 |  |  |  |
| 4 | MZ268600.1 | 21923 | 411 | T>A | AAT>AAA | p.Asn137Lys | SM -14 | India/Gandhinagar | 24-05-2021 | 25-11-2020 |
|  |  | 23353 | 1841 | A>G | GAT>GGT | p.Asp614Gly | SM-39 |  |  |  |
|  |  | 24298 | 2786 | G>T | AGT>ATT | p.Ser929Ile | SM-55 |  |  |  |
| 5 | MZ268601.1 | 21540 | 28 | C>T | CTA>TTA | p.Leu10Leu | SM -01 | India/Gandhinagar | 24-05-2021 | 24-11-2020 |
|  |  | 23353 | 1841 | A>G | GAT>GGT | p.Asp614Gly | SM-39 |  |  |  |
| 6 | MZ268602.1 | 23353 | 1841 | A>G | GAT>GGT | p.Asp614Gly | SM-39 | India/Gandhinagar | 24-05-2021 | 24-11-2020 |
|  |  | 23554 | 2042 | C>T | CCT>CAT | p.Pro681His | SM-46 |  |  |  |
| 7 | MZ268603.1 | 23353 | 1841 | A>G | GAT>GGT | p.Asp614Gly | SM-39 | India/Gandhinagar | 24-05-2021 | 24-11-2020 |
|  |  | 23554 | 2042 | C>T | CCT>CAT | p.Pro681His | SM-46 |  |  |  |
| 8 | MZ268619.1 | 23353 | 1841 | A>G | GAT>GGT | p.Asp614Gly | SM-39 | India/Gandhinagar | 24-05-2021 | 24-11-2020 |
| 9 | MZ268620.1 | 23353 | 1841 | A>G | GAT>GGT | p.Asp614Gly | SM-39 | India/Gandhinagar | 24-05-2021 | 24-11-2020 |
| 10 | MZ268626.1 | 22394 | 882 | C>T | GAC>GAT | p.Asp294Asp | SM -26 | India/Gandhinagar | 24-05-2021 | 24-11-2020 |
|  |  | 23353 | 1841 | A>G | GAT>GGT | p.Asp614Gly | SM-39 |  |  |  |
| 11 | MZ268627.1 | 22394 | 882 | C>T | GAC>GAT | p.Asp294Asp | SM -26 | India/Gandhinagar | 24-05-2021 | 24-11-2020 |
|  |  | 23353 | 1841 | A>G | GAT>GGT | p.Asp614Gly | SM-39 |  |  |  |
| 12 | MZ267529.1 | 23353 | 1841 | A>G | GAT>GGT | p.Asp614Gly | SM-39 | India/Gandhinagar | 23-05-2021 | 24-11-2020 |
|  |  | 23537 | 2025 | G>C | CAG>CAC | p.Gln675His | SM-43 |  |  |  |
|  |  | 24045 | 2533 | G>T | GCT>TCT | p.Ala845Ser | SM-51 |  |  |  |
| 13 | MZ266543.1 | 23353 | 1841 | A>G | GAT>GGT | p.Asp614Gly | SM-39 | India/Gandhinagar | 22-05-2021 | 11-12-2020 |
|  |  | 244416 | 2904 | C>T | TCC>TCT | p.Ser968Ser | SM-56 |  |  |  |
| 14 | MW595915 | 23362 | 1841 | A>G | GAT>GGT | p.Asp614Gly | SM-39 | India/Gandhinagar/Ahmedabad | 12-02-2021 | 10-11-2020 |
| 15 | MW242670.1 | 22402 | 882 | C>T | GAC>GAT | p.Asp294Asp | SM -26 | India/Gandhinagar/Go dhra | 11-11-2020 | 18-07-2020 |
|  |  | 23361 | 1841 | A>G | GAT>GGT | p.Asp614Gly | SM-39 |  |  |  |
|  |  | 23955 | 2435 | C>T | CCA>CTA | p.Pro812Leu | SM-50 |  |  |  |
| 16 | MW242692.1 | 23361 | 1841 | A>G | GAT>GGT | p.Asp614Gly | SM-39 | India/Gandhinagar/Sutrapada | 11-11-2020 | 14-07-2020 |
| 17 | MW242704.1 | 23361 | 1841 | A>G | GAT>GGT | p.Asp614Gly | SM-39 | India/Gandhinagar/Sutrapada | 11-11-2020 | 14-07-2020 |
| 18 | MW368700.1 | 22402 | 882 | C>T | GAC>GAT | p.Asp294Asp | SM -26 | India/Gandhinagar/Jamhambhaliya | 16-12-2020 | 11-07-2020 |
|  |  | 23361 | 1841 | A>G | GAT>GGT | p.Asp614Gly | SM-39 |  |  |  |
|  |  | 23955 | 2435 | C>T | CCA>CTA | p.Pro812Leu | SM-50 |  |  |  |
|  |  | 24792 | 3272 | G>T | CGT>CTT | p.Arg1091Leu | SM-60 |  |  |  |
| 19 | MW242962.1 | 23361 | 1841 | A>G | GAT>GGT | p.Asp614Gly | SM-39 | India/Gandhinagar/Junagadh | 12-11-2020 | 15-07-2020 |
| 20 | MW242706.1 | 23361 | 1841 | A>G | GAT>GGT | p.Asp614Gly | SM-39 | India/Gandhinagar/Veraval | 11-11-2020 | 14-07-2020 |
|  |  | 25046 | 3526 | G>T | GTT>TTT | p.Val1176Phe | SM-67 |  |  |  |
|  |  | 23955 | 2435 | C>T | CCA>CTA | p.Pro812Leu | SM-50 |  |  |  |
| 21 | MW242778.1 | 22402 | 882 | C>T | GAC>GAT | p.Asp294Asp | SM -26 | India/Gandhinagar/Talala | 11-11-2020 | 14-07-2020 |
|  |  | 23361 | 1841 | A>G | GAT>GGT | p.Asp614Gly | SM-39 |  |  |  |
|  |  | 23955 | 2435 | C>T | CCA>CTA | p.Pro812Leu | SM-50 |  |  |  |
| 22 | MZ269095.1 | 22251 | 739 | A>G | AGT>GGT | p.Ser247Val | SM-23 | India/Gandhinagar | 24-05-2021 | 17-12-2020 |
|  |  | 23353 | 1841 | A>G | GAT>GGT | p.Asp614Gly | SM-39 |  |  |  |
| 23 | MZ269100.1 | 22251 | 739 | A>G | AGT>GGT | p.Ser247Val | SM-23 | India/Gandhinagar | 24-05-2021 | 17-12-2020 |
|  |  | 22394 | 882 | C>T | GAC>GAT | p.Asp294Asp | SM -26 |  |  |  |
|  |  | 23353 | 1841 | A>G | GAT>GGT | p.Asp614Gly | SM-39 |  |  |  |
| 24 | MZ269224.1 | 23353 | 1841 | A>G | GAT>GGT | p.Asp614Gly | SM-39 | India/Gandhinagar | 24-05-2021 | 01-04-2021 |
|  |  | 23554 | 2042 | C>A | CCT>CAT | p.Pro681His | SM-46 |  |  |  |
|  |  | 24080 | 2568 | C>T | AAC>AAT | p.Asn856Asn | SM-52 |  |  |  |
| 25 | MZ269147.1 | 21580 | 68 | A>G | CAA>CGA | p.Gln23Arg | SM-05 | India/Gandhinagar | 24-05-2021 | 23-12-2020 |
|  |  | 23353 | 1841 | A>G | GAT>GGT | p.Asp614Gly | SM-39 |  |  |  |
| 26 | MT664161.1 | 21682 | 162 | G>T | TTG>TTT | p.Leu54Phe | SM-10 | India/Gandhinagar/SURAT | 25-06-2020 | 11-06-2020 |
|  |  | 22402 | 882 | C>T | GAC>GAT | p.Asp294Asp | SM -26 |  |  |  |
|  |  | 23361 | 1841 | A>G | GAT>GGT | p.Asp614Gly | SM-39 |  |  |  |
| 27 | MT664201.1 | 22402 | 882 | C>T | GAC>GAT | p.Asp294Asp | SM -26 | India/Gandhinagar/SURAT | 25-06-2020 | 11-06-2020 |
| 28 | MT664209.1 | 23361 | 1841 | A>G | GAT>GGT | p.Asp614Gly | SM-39 | India/Gandhinagar/SURAT | 25-06-2020 | 11-06-2020 |
| 29 | MT664729.1 | 21682 | 162 | G>T | TTG>TTT | p.Leu54Phe | SM-10 | India/Gandhinagar/SURAT | 25-06-2020 | 11-06-2020 |
|  |  | 22402 | 882 | C>T | GAC>GAT | p.Asp294Asp | SM -26 |  |  |  |
|  |  | 23361 | 1841 | A>G | GAT>GGT | p.Asp614Gly | SM-39 |  |  |  |
| 30 | MW595917 | 22838 | 1317 | C>A | AAC>AAA | p.Asn439Lys | SM-32 | India/Gandhinagar/DIU | 12-02-2021 | 22-12-2020 |
|  |  | 23362 | 1841 | A>G | GAT>GGT | p.Asp614Gly | SM-39 |  |  |  |
|  |  | 25047 | 3526 | G>T | GTT>TTT | p.Val1176Phe | SM-67 |  |  |  |
| 31 | MZ292133.1 | 22177 | 665 | C>T | GCT>GTT | p.Ala222Val | SM -22 | India/Gandhinagar | 26-05-2021 | 02-01-2021 |
|  |  | 22394 | 882 | C>T | GAC>GAT | p.Asp294Asp | SM -26 |  |  |  |
| 32 | MZ277281.1 | 412 | 412 | G>T | GAT>TAT | p.Asp138Tyr | SM-15 | Iran/ | 25-05-2021 | 10-10-2020 |
|  |  | 1250 | 1250 | A>C | AAG>ACG | p.Lys417Thr | SM-31 |  |  |  |
|  |  | 1430 | 1430 | G>A | AGC>AAC | p.Ser477Asx | SM-35 |  |  |  |
|  |  | 1450 | 1450 | G>A | GAA>AAA | p.Glu484Lys | SM-36 |  |  |  |
|  |  | 1501 | 1501 | A>T | AAT>TAT | p.Asn501Tyr | SM-37 |  |  |  |
|  |  | 1841 | 1841 | A>G | GAT>GGT | p.Asp614Gly | SM-39 |  |  |  |
| 33 | MW737421.1 | 24910 | 3386 | T>A | GTA>GAA | p.Val1129Glu | SM-66 | Iran/ | 13-03-2021 | 11-02-2020 |
|  |  | 23501 | 1977 | A>T | TCA>TCT | p.Ser659Ser | SM-41 |  |  |  |
|  |  | 24841 | 3317 | A>C | CAA>CCA | p.Gln1106Pro | SM-62 |  |  |  |
| 34 | MW672353.1 | 21746 | 238 | G>T | GAT>TAT | p.Asp80Tyr | SM-13 | Austria | 27-02-2021 | Feb-21 |
|  |  | 22825 | 1317 | C>A | AAC>AAA | p.Asn439Lys | SM-32 |  |  |  |
|  |  | 23349 | 1841 | A>G | GAT>GGT | p.Asp614Gly | SM-39 |  |  |  |
|  |  | 25153 | 3645 | C>T | TAC>TAT | p.Tyr1215Tyr | SM-68 |  |  |  |
| 35 | MW672354.1 | 21711-21716 | 203TO208DEL | TACATG - DELETION | TAC ATGdel |  | SM -12 | Austria | 27-02-2021 | Feb-21 |
|  |  | 22073 | 565 | C>T | CTT>TTT | p.Leu189Phe | SM-20 |  |  |  |
|  |  | 22825 | 1317 | C>A | AAC>AAA | p.Asn439Lys | SM-32 |  |  |  |
|  |  | 23349 | 1841 | A>G | GAT>GGT | p.Asp614Gly | SM-39 |  |  |  |
|  |  | 23822 | 2314 | G>A | GTT>ATT | p.Val772Ile | SM-49 |  |  |  |
|  |  | 24860 | 3348 | T>C | ACT>ACC | p.Thr1116Thr | SM-63 |  |  |  |
| 36 | MW672355.1 | 21711-21716 | 203TO208DEL | TACATG - DELETION | TAC ATGdel |  | SM -12 | Austria | 27-02-2021 | Feb-21 |
|  |  | 22073 | 565 | C>T | CTT>TTT | p.Leu189Phe | SM-20 |  |  |  |
|  |  | 22825 | 1317 | C>A | AAC>AAA | p.Asn439Lys | SM-32 |  |  |  |
|  |  | 23349 | 1841 | A>G | GAT>GGT | p.Asp614Gly | SM-39 |  |  |  |
|  |  | 23822 | 2314 | G>A | GTT>ATT | p.Val772Ile | SM-49 |  |  |  |
|  |  | 24856 | 3348 | T>C | ACT>ACC | p.Thr1116Thr | SM-63 |  |  |  |
| 37 | MW723179<br>.1 | 21702-21707 | 203TO208<br>DEL | TACATG -<br>DELETIO<br>N | TAC ATGdel |  | SM -12 | Austria | 10-03-2021 | 17-02-2021 |
|  |  | 21930-21932 | 431TO433 | ATT-DELETIO<br>N | ATTdel |  | SM-16 |  |  |  |
|  |  | 21962 | 463 | A>G | AGT>GGT | p.Ser155Gly | SM-19 |  |  |  |
|  |  | 23000 | 1501 | A>T | AAT>TAT | p.Asn501Tyr | SM-37 |  |  |  |
|  |  | 23208 | 1709 | C>A | GCT>GAT | p.Ala570Asp | SM-38 |  |  |  |
|  |  | 23340 | 1841 | A>G | GAT>GGT | p.Asp614Gly | SM-39 |  |  |  |
|  |  | 23541 | 2042 | C>A | CCT>CAT | p.Pro681His | SM-46 |  |  |  |
|  |  | 23646 | 2147 | C>T | ACA>ATA | p.Thr716Ile | SM-48 |  |  |  |
|  |  | 24443 | 2944 | T>G | TCA>GCA | p.Ser982Ala | SM-57 |  |  |  |
|  |  | 24851 | 3352 | G>C | GAC>CAC | p.Asp1118His | SM-64 |  |  |  |
| 38 | MW723183<br>.1 | 21746 | 238 | G>T | GAT>TAT | p.Asp80Tyr | SM-13 | Austria | 10-03-2021 | 19-02-2021 |
|  |  | 22825 | 1317 | C>A | AAC>AAA | p.Asn439Lys | SM-32 |  |  |  |
|  |  | 23349 | 1841 | A>G | GAT>GGT | p.Asp614Gly | SM-39 |  |  |  |
| 39 | MT318827.<br>1 | 23403 | 1841 | A>G | GAT>GGT | p.Asp614Gly | SM-39 | Germany | 04-05-2020 | 19-03-2020 |
| 40 | MT358639.<br>1 | 23103 | 1841 | A>G | GAT>GGT | p.Asp614Gly | SM-39 | Germany | 14-05-2020 | Feb-20 |
| 41 | MT270102.<br>1 | 23374 | 1841 | A>G | GAT>GGT | p.Asp614Gly | SM-39 | Germany:<br>Bavaria | 19-05-2020 | 28-01-2020 |
| 42 | MT582454.<br>1 | 21573 | 65 | C>T | ACT>ATT | p.Thr22Ile | SM-04 | Germany:<br>Dusseldorf | 09-06-2020 | 21-03-2020 |
| 43 | MT582497.<br>1 |  |  |  |  |  |  | Germany:<br>Heinsberg | 09-06-2020 | 26-02-2020 |
| 44 | MW397523<br>.1 | 22223 | 665 | C>T | GCT>GTT | p.Ala222Val | SM -22 | Italy | 215-21 | 04-11-2020 |
|  |  | 23399 | 1841 | A>G | GAT>GGT | p.Asp614Gly | SM-39 |  |  |  |
|  |  | 23583 | 2025 | G>T | CAG>CAT | p.Gln675His | SM-42 |  |  |  |
| 45 | MZ054387.<br>1 | 22195 | 665 | C>T | GCT>GTT | p.Ala222Val | SM -22 | Italy | 04-05-2021 | 15-02-2021 |
|  |  | 22314 | 784 | G>T | GCT>GTT | p.Ala262Val | SM -24 |  |  |  |
|  |  | 22345 | 815 | C>T | CCT>CTT | p.Pro272Leu | SM -25 |  |  |  |
|  |  | 23371 | 1841 | A>G | GAT>GGT | p.Asp614Gly | SM-39 |  |  |  |
| 46 | MW852494<br>.1 | 21701 | 155 | A>G | CAG>CGG | p.Gln52Arg | SM -09 | Italy | 02-04-2021 | 15-03-2021 |
|  |  | 21746 | 200 | C>T | GCT>GTT | p.Ala67Val | SM-11 |  |  |  |
|  |  | 21749-21754 | 203TO208<br>DEL | TACATG -<br>DELETIO<br>N | TAC ATGdel |  | SM-12 |  |  |  |
|  |  | 21977-21979 | 431TO433 | ATT-DELETIO<br>N | ATTdel |  | SM-16 |  |  |  |
|  |  | 22004 | 458 | T>C | ATG>ACG | p.Met153Thr | SM-18 |  |  |  |
|  |  | 22602 | 1056 | T>A | GCT>GCA | p.Ala352Ala | SM-28 |  |  |  |
|  |  | 22747 | 1201 | G>T | GTA>TTA | p.Val401Leu | SM-30 |  |  |  |
|  |  | 23387 | 1841 | A>G | GAT>GGT | p.Asp614Gly | SM-39 |  |  |  |
|  |  | 23577 | 2031 | G>C | CAG>CAC | p.Gln677His | SM-45 |  |  |  |
|  |  | 24208 | 2662 | T>C | TTT>CTT | p.Phe888Leu | SM-54 |  |  |  |
|  |  | 24732 | 3186 | C>T | TTC>TTT | p.Phe1062Phe | SM-59 |  |  |  |
|  |  | 25367 | 3821 | A>G | TAA>TGA | p.*1274* | SM-70 |  |  |  |
| 47 | MW854297<br>.1 | 21749-21754 | 203TO208<br>DEL | TACATG -<br>DELETIO<br>N | TAC ATGdel |  | SM -12 | Italy | 02-04-2021 | 10-03-2021 |
|  |  | 21977-21979 | 431TO433 | ATT-DELETIO<br>N | ATTdel |  | SM-16 |  |  |  |
|  |  | 22542 | 906 | G>T | ACG>ACT | p.Thr302Thr | SM-27 |  |  |  |
|  |  | 23047 | 1501 | A>T | AAT>TAT | p.Asn501Tyr | SM-37 |  |  |  |
|  |  | 23255 | 1709 | C>A | GCT>GAT | p.Ala570Asp | SM-38 |  |  |  |
|  |  | 23387 | 1841 | A>G | GAT>GGT | p.Asp614Gly | SM-39 |  |  |  |
|  |  | 23588 | 2042 | C>A | CCT>CAT | p.Pro681His | SM-46 |  |  |  |
|  |  | 23693 | 2147 | C>T | ACA>ATA | p.Thr716Ile | SM-48 |  |  |  |
|  |  | 24490 | 2944 | T>G | TCA>GCA | p.Ser982Ala | SM-57 |  |  |  |
|  |  | 24898 | 3352 | G>C | GAC>CAC | p.Asp1118His | SM-64 |  |  |  |
| 48 | MW786740<br>.1 | 21621 | 84 | C>T | TAC>TAT | p.Tyr28Tyr | SM -07 | Italy | 22-03-2021 | 22-03-2021 |
|  |  | 21968 | 431 | A>T | TAT>TTT | p.Tyr144Phe | SM-17 |  |  |  |
|  |  | 22202 | 665 | C>T | GCT>GTT | p.Ala222Val | SM -22 |  |  |  |
|  |  | 23378 | 1841 | A>G | GAT>GGT | p.Asp614Gly | SM-39 |  |  |  |
| 49 | MW332227<br>.1 | 23373 | 1841 | A>G | GAT>GGT | p.Asp614Gly | SM-39 | Russia:<br>Moscow | 06-12-2020 | 18-03-2020 |
| 50 | MW332226<br>.1 | 23377 | 1841 | A>G | GAT>GGT | p.Asp614Gly | SM-39 | Russia:<br>Saratov | 06-12-2020 | 30-07-2020 |
| 51 | MW332228<br>.1 | 23373 | 1841 | A>G | GAT>GGT | p.Asp614Gly | SM-39 | Russia:<br>Samara | 06-12-2020 | 11-04-2020 |
| 52 | MW332240<br>.1 | 23373 | 1841 | A>G | GAT>GGT | p.Asp614Gly | SM-39 | Russia:<br>Penza | 06-12-2020 | 08-04-2020 |
| 53 | MZ277285.<br>1 | 21718-21723 | 203TO208<br>DEL | TACATG -<br>DELETIO<br>N | TAC ATGdel |  | SM -12 | USA:<br>California,<br>Los Angeles | 25-05-2021 | 06-05-2021 |
|  |  | 21946-21948 | 431TO433 | ATT-DELETIO<br>N | ATTdel |  | SM-16 |  |  |  |
|  |  | 23016 | 1501 | A>T | AAT>TAT | p.Asn501Tyr | SM-37 |  |  |  |
|  |  | 23224 | 1709 | C>A | GCT>GAT | p.Ala570Asp | SM-38 |  |  |  |
|  |  | 23356 | 1841 | A>G | GAT>GGT | p.Asp614Gly | SM-39 |  |  |  |
|  |  | 23557 | 2042 | C>A | CCT>CAT | p.Pro681His | SM-46 |  |  |  |
|  |  | 23662 | 2147 | C>T | ACA>ATA | p.Thr716Ile | SM-48 |  |  |  |
|  |  | 24459 | 2944 | T>G | TCA>GCA | p.Ser982Ala | SM-57 |  |  |  |
|  |  | 24867 | 3352 | G>C | GAC>CAC | p.Asp1118His | SM-64 |  |  |  |
| 54 | MZ277986.1 | 21551 | 52 | C>T | CTT>TTT | p.Leu18Phe | SM-02 | USA: Texas | 25-05-2021 | 07-05-2021 |
|  |  | 21558 | 59 | C>A | ACC>AAC | p.Thr20Asn | SM-03 |  |  |  |
|  |  | 21575 | 76 | C>T | CCT>TCT | p.Pro26Ser | SM-06 |  |  |  |
|  |  | 21911 | 412 | G>T | GAT>TAT | p.Asp138Tyr | SM-15 |  |  |  |
|  |  | 22069 | 570 | G>T | AGG>AGT | p.Arg190Ser | SM-21 |  |  |  |
|  |  | 22749 | 1250 | A>C | AAG>ACG | p.Lys417Thr | SM-31 |  |  |  |
|  |  | 23000 | 1501 | A>T | AAT>TAT | p.Asn501Tyr | SM-37 |  |  |  |
|  |  | 23340 | 1841 | A>G | GAT>GGT | p.Asp614Gly | SM-39 |  |  |  |
|  |  | 23462 | 1963 | C>T | CAT>TAT | p.His655Tyr | SM-40 |  |  |  |
|  |  | 24579 | 3080 | C>T | ACT>ATT | p.Thr1027Ile | SM-58 |  |  |  |
|  |  | 25025 | 3526 | G>T | GTT>TTT | p.Val1176Phe | SM-67 |  |  |  |
| 55 | MZ277985.1 | 21551 | 52 | C>T | CTT>TTT | p.Leu18Phe | SM-02 | USA: Pennsylvania | 25-05-2021 | 06-05-2021 |
|  |  | 21558 | 59 | C>A | ACC>AAC | p.Thr20Asn | SM-03 |  |  |  |
|  |  | 21575 | 76 | C>T | CCT>TCT | p.Pro26Ser | SM-06 |  |  |  |
|  |  | 21911 | 412 | G>T | GAT>TAT | p.Asp138Tyr | SM-15 |  |  |  |
|  |  | 22069 | 570 | G>T | AGG>AGT | p.Arg190Ser | SM-21 |  |  |  |
|  |  | 22749 | 1250 | A>C | AAG>ACG | p.Lys417Thr | SM-31 |  |  |  |
|  |  | 23000 | 1501 | A>T | AAT>TAT | p.Asn501Tyr | SM-37 |  |  |  |
|  |  | 23340 | 1841 | A>G | GAT>GGT | p.Asp614Gly | SM-39 |  |  |  |
|  |  | 23462 | 1963 | C>T | CAT>TAT | p.His655Tyr | SM-40 |  |  |  |
|  |  | 24579 | 3080 | C>T | ACT>ATT | p.Thr1027Ile | SM-58 |  |  |  |
|  |  | 25025 | 3526 | G>T | GTT>TTT | p.Val1176Phe | SM-67 |  |  |  |
| 56 | MZ277987.1 | 21551 | 52 | C>T | CTT>TTT | p.Leu18Phe | SM-02 | USA: Missouri | 25-05-2021 | 07-05-2021 |
|  |  | 21558 | 59 | C>A | ACC>AAC | p.Thr20Asn | SM-03 |  |  |  |
|  |  | 21575 | 76 | C>T | CCT>TCT | p.Pro26Ser | SM-06 |  |  |  |
|  |  | 21911 | 412 | G>T | GAT>TAT | p.Asp138Tyr | SM-15 |  |  |  |
|  |  | 22069 | 570 | G>T | AGG>AGT | p.Arg190Ser | SM-21 |  |  |  |
|  |  | 22614 | 1115 | C>T | GCA>GTA | p.Ala372Val | SM-29 |  |  |  |
|  |  | 22749 | 1250 | A>C | AAG>ACG | p.Lys417Thr | SM-31 |  |  |  |
|  |  | 23000 | 1501 | A>T | AAT>TAT | p.Asn501Tyr | SM-37 |  |  |  |
|  |  | 23340 | 1841 | A>G | GAT>GGT | p.Asp614Gly | SM-39 |  |  |  |
|  |  | 23462 | 1963 | C>T | CAT>TAT | p.His655Tyr | SM-40 |  |  |  |
|  |  | 23562 | 2063 | C>T | GCT>GTT | p.Ala688Val | SM-47 |  |  |  |
|  |  | 24579 | 3080 | C>T | ACT>ATT | p.Thr1027Ile | SM-58 |  |  |  |
|  |  | 25025 | 3526 | G>T | GTT>TTT | p.Val1176Phe | SM-67 |  |  |  |
| 57 | MZ277988.1 | 21551 | 52 | C>T | CTT>TTT | p.Leu18Phe | SM-02 | USA: Ohio | 25-05-2021 | 07-05-2021 |
|  |  | 21558 | 59 | C>A | ACC>AAC | p.Thr20Asn | SM-03 |  |  |  |
|  |  | 21575 | 76 | C>T | CCT>TCT | p.Pro26Ser | SM-06 |  |  |  |
|  |  | 21911 | 412 | G>T | GAT>TAT | p.AsP138Tyr | SM-15 |  |  |  |
|  |  | 22069 | 570 | G>T | AGG>AGT | p.Arg190Ser | SM-21 |  |  |  |
|  |  | 22749 | 1250 | A>C | AAG>ACG | p.Lys417Thr | SM-31 |  |  |  |
|  |  | 23000 | 1501 | A>T | AAT>TAT | p.Asn501Tyr | SM-37 |  |  |  |
|  |  | 23340 | 1841 | A>G | GAT>GGT | p.Asp614Gly | SM-39 |  |  |  |
|  |  | 23462 | 1963 | C>T | CAT>TAT | p.His655Tyr | SM-40 |  |  |  |
|  |  | 23562 | 2063 | C>T | GCT>GTT | p.Ala688Val | SM-47 |  |  |  |
|  |  | 23934 | 2435 | C>T | CCA>CTA | p.Pro812Leu | SM-50 |  |  |  |
|  |  | 24579 | 3080 | C>T | ACT>ATT | p.Thr1027Ile | SM-58 |  |  |  |
|  |  | 24772 | 3273 | T>C | CGT>CGC | p.Arg1091Arg | SM-61 |  |  |  |
|  |  | 25025 | 3526 | G>T | GTT>TTT | p.Val1176Phe | SM-67 |  |  |  |
| 58 | MW691151.1 | 22437 | 882 | C>T | GAC>GAT | p.Asp294Asp | SM -26 | China:<br>Hong Kong | 23-03-2021 | 27-12-2020 |
|  |  | 23396 | 1841 | A>G | GAT>GGT | p.Asp614Gly | SM-39 |  |  |  |
|  |  | 24168 | 2613 | T>C | GCT>GCC | p.Ala871Ala | SM-53 |  |  |  |
| 59 | MW301121.1 | 23397 | 1841 | A>G | GAT>GGT | p.Asp614Gly | SM-39 | China:<br>Chengdu,Sichuan province | 28-11-2020 | 10-03-2020 |
| 60 | MW011762<br>.1 |  |  |  |  |  |  | China:<br>Hunan,Huai<br>hua | 16-09-2020 | 17-01-2020 |
| 61 | MW011765<br>.1 |  |  |  |  |  |  | China:<br>Hunan,Shao<br>yang | 16-09-2020 | 27-01-2020 |
| 62 | MW011763<br>.1 |  |  |  |  |  |  | China:<br>Hunan,Huai<br>hua | 16-09-2020 | 25-01-2020 |
| 63 | LC632047.<br>1 | 23391 | 1841 | A>G | GAT>GGT | p.Asp614Gly | SM-39 | Japan | 14-05-2021 | Sep-20 |
| 64 | LC630936.<br>1 | 23403 | 1841 | A>G | GAT>GGT | p.Asp614Gly | SM-39 | Japan:<br>Kanagawa,<br>Sagamihara | 11-05-2021 | 06-04-2020 |
| 65 | LC632048.<br>1 | 23402 | 1841 | A>G | GAT>GGT | p.Asp614Gly | SM-39 | Japan | 14-05-2021 | Sep-20 |
| 66 | LC632049.<br>1 | 23388 | 1841 | A>G | GAT>GGT | p.Asp614Gly | SM-39 | Japan | 14-05-2021 | Sep-20 |
| 67 | LC623934.<br>1 | 23401 | 1841 | A>G | GAT>GGT | p.Asp614Gly | SM-39 | Japan | 22-04-2021 | Sep-20 |
| 68 | MT478018.<br>1 | 23396 | 1841 | A>G | GAT>GGT | p.Asp614Gly | SM-39 | Turkey | 19-05-2021 | Mar-20 |
| 69 | MW308549<br>.1 | 23371 | 1841 | A>G | GAT>GGT | p.Asp614Gly | SM-39 | Turkey | 12-03-2021 | 18-07-2020 |
| 70 | MW320689<br>.1 | 23371 | 1841 | A>G | GAT>GGT | p.Asp614Gly | SM-39 | Turkey | 12-03-2021 | 18-07-2020 |
| 71 | MW340910<br>.1 | 23371 | 1841 | A>G | GAT>GGT | p.Asp614Gly | SM-39 | Turkey | 12-03-2021 | 18-07-2020 |
| 72 | NC_045512<br>* |  |  |  |  |  |  | CHINA | 18-JUL-2020 | Dec-19 |

<sup>a</sup>Nucleotide position of the mutation in the corresponding GenBank sequence; <sup>b</sup>Nucleotide position in the Surface Glycoprotein gene considering nucleotide A of the start codon as 1; <sup>c</sup>Change in nucleotide; <sup>d</sup>Change in Amino Acid; <sup>e</sup>Country/city from where the sequence is deposited in GenBank; <sup>f</sup>Date of release of sequence by GeneBank; <sup>g</sup>Date of collection of clinical sample used for generating the sequence.
\*Reference sequence

A set of 105 human miRNA seed location sites were detected with hit-score ranging between 92 and 50. Out of them, 11 targets had a prediction score of > 80 and hence were most likely to be real. [23] In the reference gene (NC_045512), at position 845 there is amino acid Alanine (GCT). A miRNA seed location encompassing this position was found to be significant since it was the binding site of at least 7 different human miRNAs with target score ranging from 87 to 84 (Figure 1). In our study population, this mutation (A845S) was detected in only one of the sequences submitted from India (Table 2).

**Figure 1:**
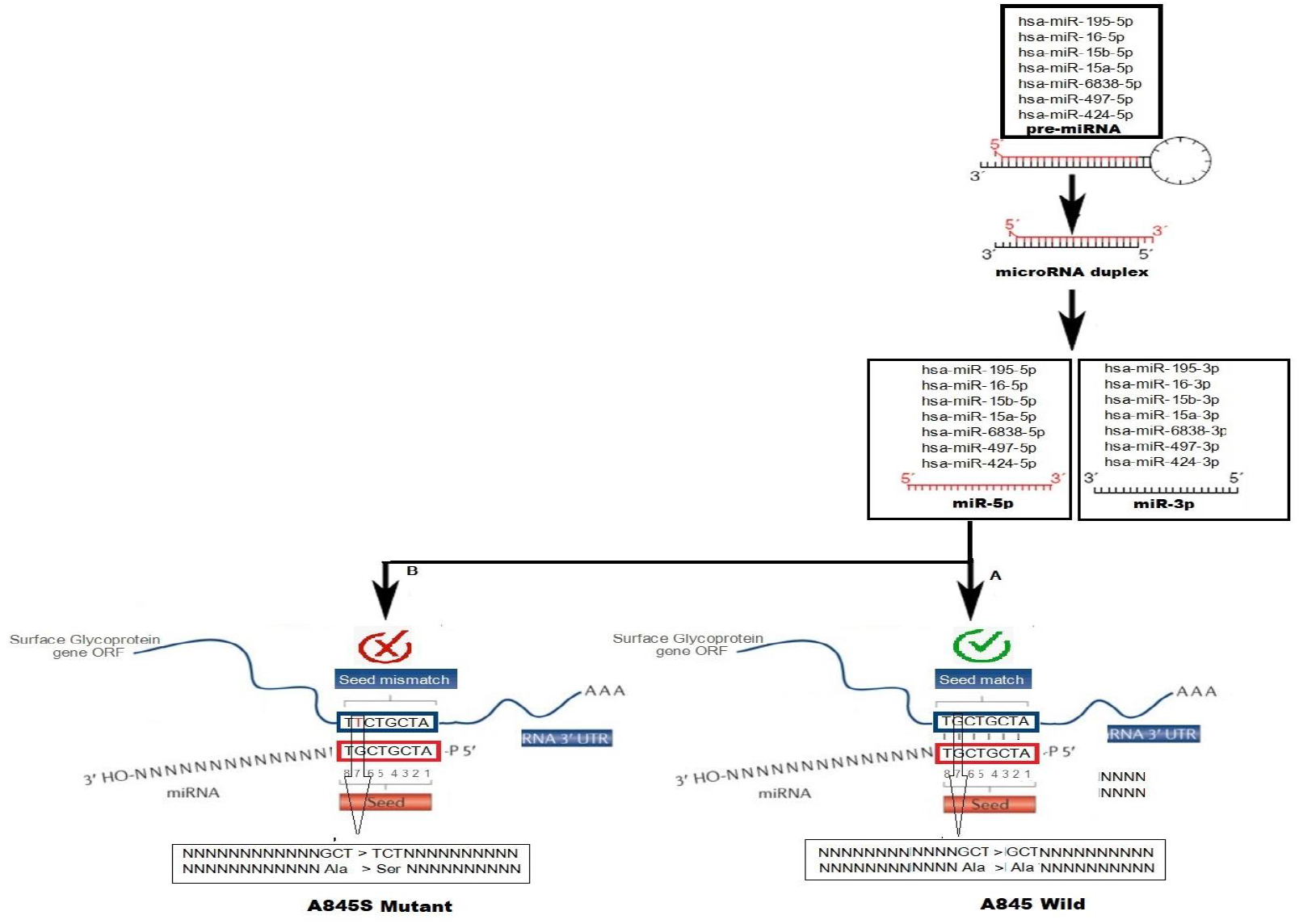
A set of 7 human miRNA (hsa-miR-195-5p, hsa-miR-16-5p, hsa-miR-15b-5p, hsa-miR-15a-5p, hsa-miR-6838-5p, hsa-miR-497-5p, hsa-miR-424-5p) with sequence complementarity encompassing amino acid position 845 (A845) in A: reference (wild) gene and B: With mutation (A845S) in an Indian sequence where the sequence complementarity is lost (B).

**Table 2:** Human microRNA along with their seed location within the Surface Glycoprotein (S) gene and mutations observed within the region.

| miRDB<sup>a</sup> Target Score | miRNA<sup>b</sup> | Nucleotide location<sup>c</sup> | Laboratory mutation code | Mutation | Country |
| --- | --- | --- | --- | --- | --- |
| 89 | miR-511-3p | 3814 to 3821 | SM-70 | *1274* | Italy |
| 87 | miR-195-5p | 2532 to 2539 | SM-51 | A845S | INDIA |
| 87 | miR-16-5p | 2532 to 2539 | SM-51 | A845S | INDIA |
| 87 | miR-15b-5p | 2532 to 2539 | SM-51 | A845S | INDIA |
| 87 | miR-15a-5p | 2532 to 2539 | SM-51 | A845S | INDIA |
| 85 | miR-219a-1-3p | 1971 to 1977 | SM-41 | S659S | Iran |
| 84 | miR-6838-5p | 2532 to 2539 | SM-51 | A845S | INDIA |
| 84 | miR-497-5p | 2532 to 2539 | SM-51 | A845S | INDIA |
| 84 | miR-424-5p | 2532 to 2539 | SM-51 | A845S | INDIA |
| 81 | miR-20b-3p | 3346 to 3353 | SM-63 | T1116T | Austria |
|  |  |  | SM-64 | D1118H | Austria, Italy, USA |

<sup>a</sup>http://mirdb.org/; <sup>b</sup>Human microRNA (miRNA) with seed sequence within the coding region of Surface Glycoprotein (S) gene; <sup>c</sup>Region within the S gene encompassing the seed sequence.

A synonymous mutation encompassing the stop codon but retaining the same was detected in one of the S gene sequences submitted from Italy. The synonymous nature of this mutation altered an Ochre stop codon (UAA) to an Opal one (UGA) (Figure 2). This mutation is predicted to have no effect on the protein synthesis mechanism of the strain but is anticipated to erase a potential human miRNA (miR-511-3p) seed sequence complementarity from this location. Similar synonymous mutation was encountered in two other locations within the S gene. These are amino acid position 659 and 1116 where an SNP was detected but the amino acid coded remained to be Serine and Threonine, thereby altering the seed sequence for miR-219a-1-3p and miR-20b-3p in S gene sequences reported from Iran and Austria respectively (Figure 3 and Figure 4). miR-20b-3p had yet another seed location at amino acid position 1118 within the S gene which was found to be mutated (D1118H) in the Austrian sequence along with two others from Italy and USA (Figure 4).

**Figure 2:**
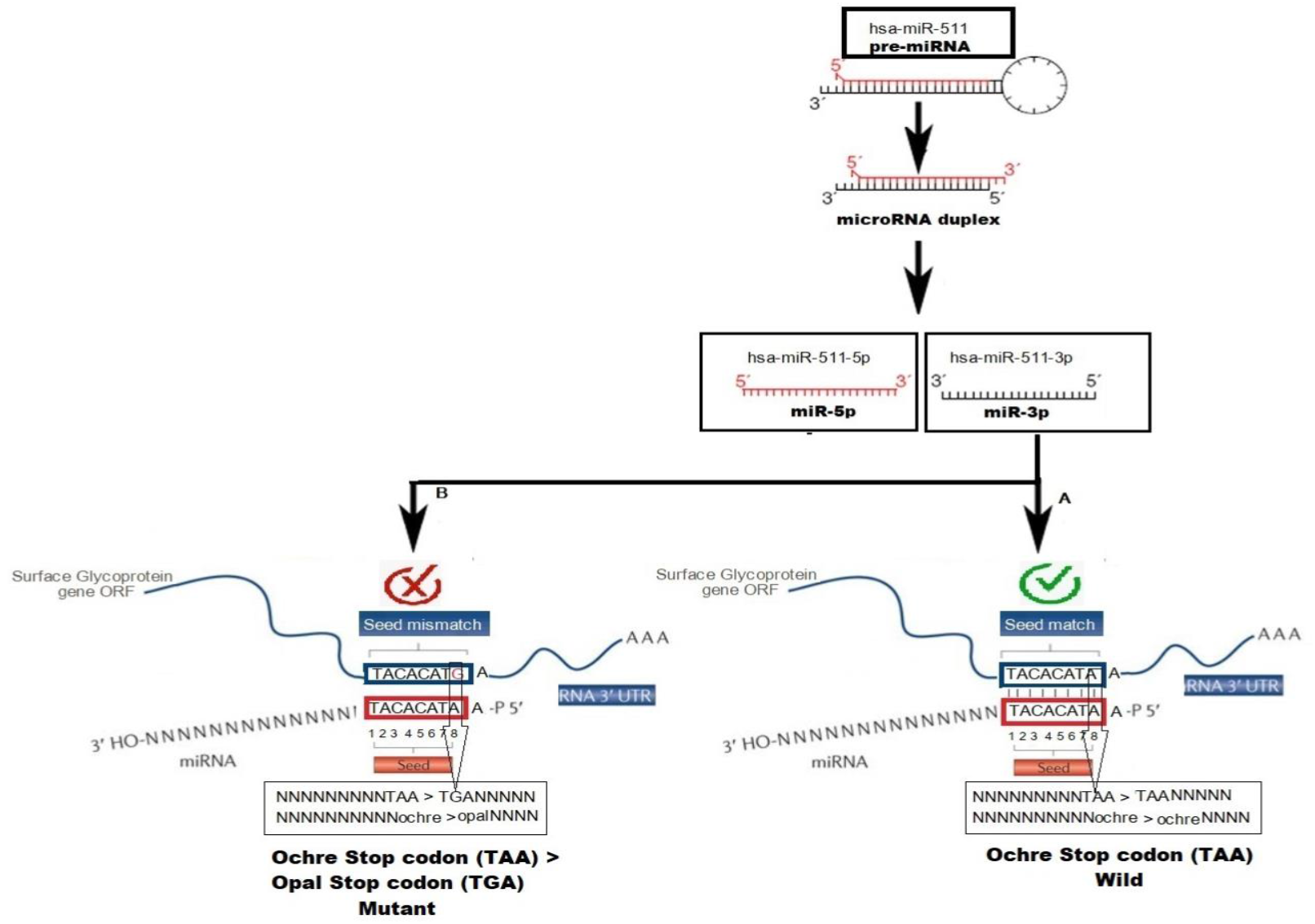
Mutation in S gene at amino acid position 1274 retaining the stop codon but altering a single base within, to abolish seed sequence complementarity of miR-511-5p. A: Reference sequence showing Ochre stop codon (TAA) and B: Sequence with Accession No. MW852494.1 showing an Opal stop codon (TGA).

**Figure 3:**
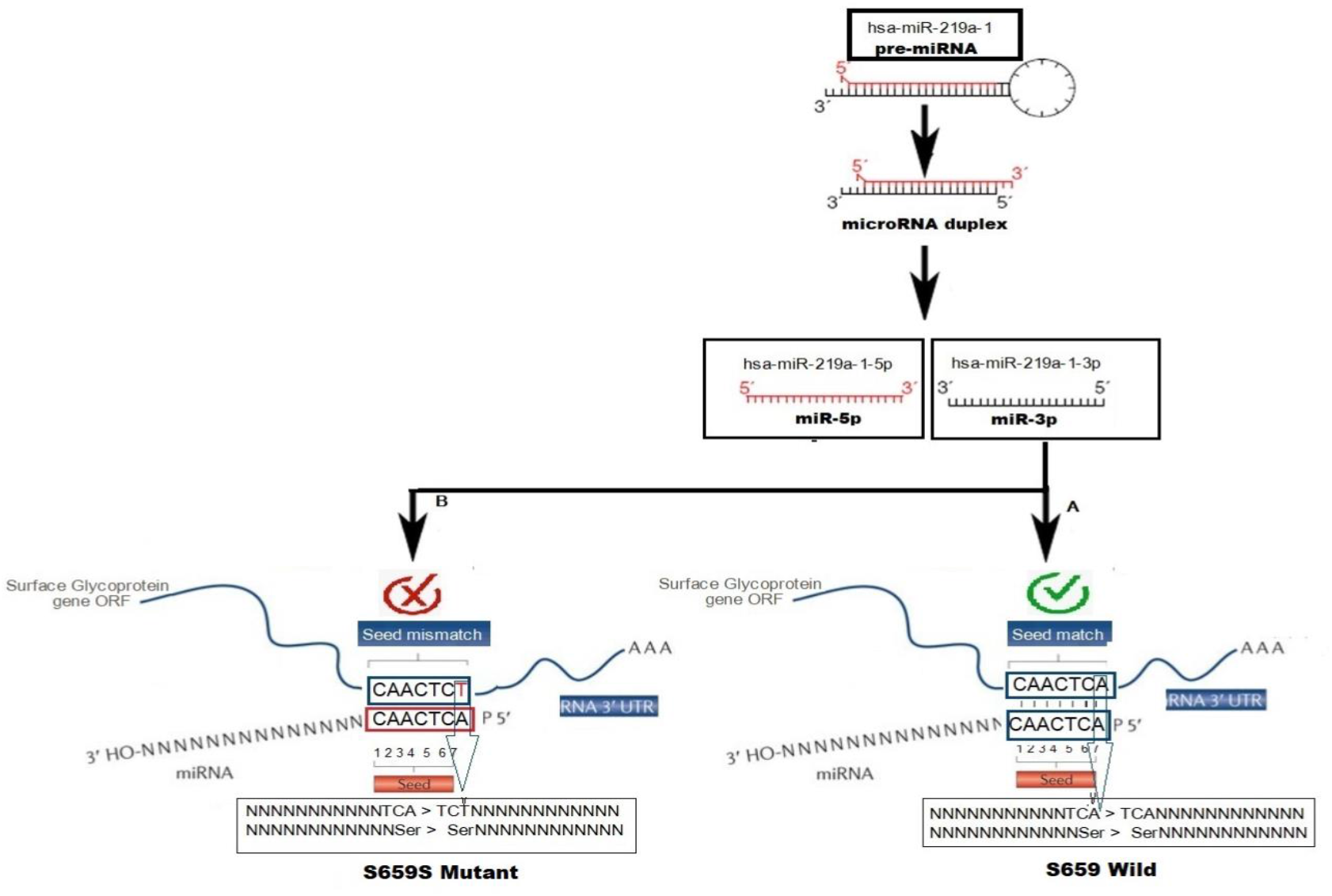
miRNA miR-219a-1-3p losing sequence complementarity due to a synonymous mutation at amino acid position 659 (S659S). A: Reference sequence (Accession No.NC_045512) B: Mutated Sequence (Accession no.MW737421.1).

**Figure 4:**
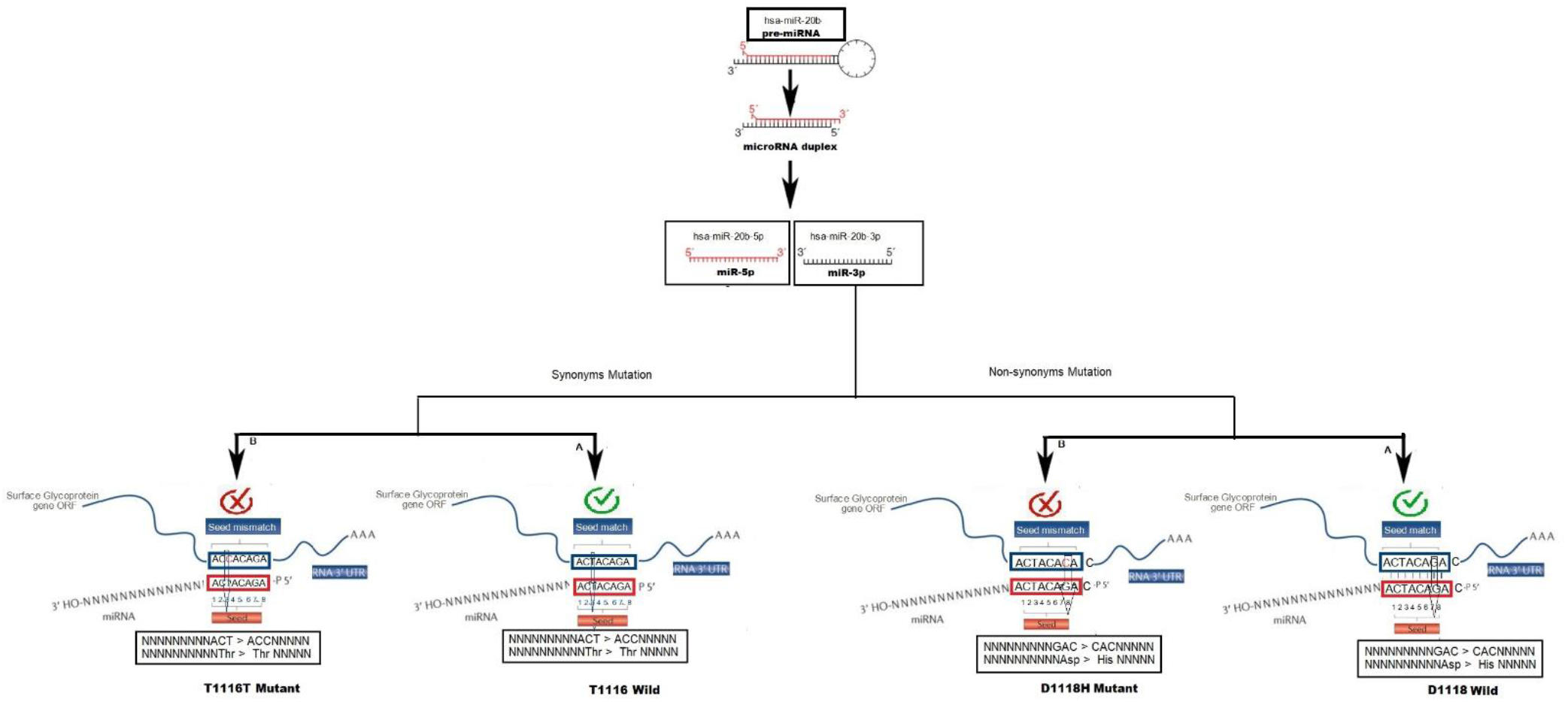
miRNA miR-20b-3p losing two sees sequence complementarity at positions 1116 (T1116T-a synonymous mutation) and 1118 (D1118H - a non-synonymous mutation). Left: 1116 Mutation; Right: 1118 mutation; A: Reference sequence (Accession No. NC_045512); B: mutated sequence (For position 1116: Accession No. MW672354.1,MW672355.1; For position 1118: MW723179.1,MW854297.1, MZ277285.1).

## Discussion

Since early 2020, the COVID-19 epidemic has changed social behaviour as much as national economies by killing a massive population of the world. [24] Our past experience with severe acute respiratory syndrome coronavirus (SARS-CoV) and Middle East respiratory syndrome (MERS) indicated that the pathogen jumped from animals to humans. [25] Given the similarity between SARS-CoV and the current SARS-CoV-2, a similar mode of transmission seems logical. [26] With the virus spreading fast and furious in India that culminated in so-called second wave, LDA (Latent Dirichlet Allocation) topic modelling based on social media platform data indicated fear of death, unemployment, and worry being rampant in the Indian population post the second wave of COVID-19 attack that started in April 2021. [27]

In several instances, host miRNA has been demonstrated to carry out antiviral functions by reducing the level of viral infection inside the host. A classic example is human microRNA miR-32 that anneals to the mRNA of primate foamy virus type 1 retrovirus thereby restricting the accumulation of viral RNA in human cells. [28] Similarly, viral large protein (L protein) and phosphoprotein (P protein) genes of the vesicular stomatitis virus (VSV), a negative-sense RNA virus, has been shown to be targeted by the human miR-24 and miR-93 to activate potent antiviral defence by the host. [29] Human miR-29a has been shown to attack the 3’ UTR of human immunodeficiency virus type 1 (HIV-1, a retrovirus) RNA inside the T-lymphocytes to reduce the HIV-1 viral replication and infectivity. [30] Human miR-145 directly binds and inhibits the E1 and E2 open reading frames of Human Papillomavirus (HPV) and reduces viral replication as well as expression of its late genes. [31]

Seed sequence of a microRNA is a short stretch of nucleotides and with perfect Watson-Crick complementarity with its target. This term was introduced by Lewis and co-workers while developing the TargetScan algorithm. [54] In this study, we demonstrate emergence of mutation within regions of the SARS-CoV-2 spike glycoprotein gene that are potential seed locations for several human miRNAs. We hypothesize that these mutations are events of the selection process that became statistically prominent following global outburst of infection in the form of COVID-19 waves. They provided a crucial selection advantage to the pathogen with abundant growth options and accounted for one of the several reasons for enhanced morbidity and mortality associated with COVID-19 disease worldwide.

In our analysis, 105 human miRNA with scores ranging from 50 to 92 were found to have potential seed sequence within the SARS-CoV-2 S gene (Table 1). However, a subset of miRNA with score >80 on the miRDB scale were considered for analysis because of their enhanced specificity and likelihood to be real (Table 2). [23]

A potential seed region of 8 bases (uacacaua) was detected for the human miRNA miR-511-3p in the spike gene of SARS-CoV-2. This location overlapped the Ochre stop codon (UAA) of the S gene located at the nucleotide position 25382-25384 (Figure 2); (Genbank Accession Number NC_045512). The miRNA is part of the hypothalamic miRNA group and has been shown to bind directly to ACE2 and TMPRSS2, the two human genes that play critical roles in SARS-CoV-2 pathogenesis. ACE2 is not only the entry receptor of the virus but is also known to protect the lungs from injury. [20] SARS-CoV-2 employs endosomal cysteine proteases cathepsin B and L (CatB/L) [32] and the serine protease TMPRSS2 [33,34,35] for priming of the S protein. This entails cleavage of the protein at two potential sites, Arg685/Ser686 and Arg815/Ser816. It has been demonstrated that inhibition of both these proteases is essential for blockage the entry of the virus inside the human cell. [36] Our study further strengthened the idea of inhibitory effect of this miRNA and added a dimension to its negative regulation on SARS-CoV-2 by demonstrating its *in silico* binding sites covering the stop codon region of the S glycoprotein gene. Our *in-silico* analysis demonstrated that in an isolate from Italy (Genbank accession number MW852494.1; Sample collected on 15-03-2021 and sequence released on 02-04-2021) a synonymous mutation was encountered that altered this ochre stop codon (UAA) to Opal (UGA) at position 25382-25384 thereby erasing seed location of the human miRNA miR-511-3p as mentioned above (Figure 2). This molecular event is likely to have pathogenic consequences given the fact that this miRNA has potential to bind to the 3’ untranslated regions of the wild type S gene exploiting this seed sequence and suppress translation of spike proteins for the virus.

A single SARS-CV-2 S gene sequence submitted from India (out of a total of 31) was found to encounter a mutation at amino acid position 845 that resulted in substitution of the amino acid Alanine with Serine (A845S). This mutation erased potential seed location of a set of 7 miRNAs, namely miR-195-5p, miR-16-5p, miR-15b-5p, miR-15a-5p, miR-497-5p, miR-424-5p and miR-6838-5p (Figure 1). We hypothesize that the infecting viruses with this mutation enjoyed a survival advantage by evading these host miRNAs that are active in the human system. Barring miR-6838-5p, all these miRNAs, commonly recognizing the S gene region and encompassing amino acid position A845, form part of the miR-15/107 family, also known as the miT-15/107 gene group. They are multiple in number, highly conserved and have a common and conserved seed target on the S gene and with a high miRDB score of >80. [37] They collectively anneal at location encompassing mutation A845S detected in the Indian sequence (Accession number MZ267529) generated from a clinical sample collected on 24 November, 2020.

The miR-195-5p has been shown to express in the mammalian respiratory epithelial cells. This enhances the probability of this miRNA to physically encounter SARS-CoV-2 in an infected patient and subsequent mount antiretroviral effects. [38,39,23] Similarly, the miR-16-5p, miR-15b-5p, miR-15a-5p, miR-497-5p and miR-424-5p of this group were also identified to be important in the context of SARS-CoV-2 biology. The miRNA miR-16-5p of this group along with few others (miR-21-3p, miR-195-5p, miR-3065-5p, miR-424-5p and miR-421) were identified by Nersisyan and co-workers (2020) [40] as one that had high-confidence interaction with all seven coronaviruses known till date. Specifically, miR-16-5p along with miR-21-3p were found to be the top 5% of all miRNA experimented by the authors that were expressed across all clinical samples taken. Enterovirus 71 (EV71) is known to be the causative pathogen of hand-foot-and-mouth disease (HFMD). miR-16-5p has been demonstrated to promote EV71-induced nerve cell apoptosis through activation of caspase-3. miR-16-5p also inhibited EV71 replication. CCNE1 and CCND1, two important regulators of the cell cycle play crucial roles in the suppression of EV71 replication by miR-16-5p. [41]. Hence, it can be predicted that erasing of a potential seed sequence for the miR-16-5p on the S gene of infecting SARS-CoV-2 might have provided a replicative advantage to the pathogen in the human population during the pandemic.

Authors working on Hamster Lung Tissues Infected by SARS-CoV-2 demonstrated that different miRNAs exhibited patterns of variable expression after infection. These included miR-15b-5p and miR-15a-5p along with miR-195-5p. MicroRNA miR-15b-5p and miR-195-5p demonstrated large differences in expression in the lungs indicating that they may be potential markers for a SARS-CoV-2 infection in an individual.[42]

Jafarinejad-Farsangi and co-workers (2020) [43] undertook an *in-silico* analysis and reported that miR-497-5p was part of a group of human miRNAs that had >3 binding sites on SARS-CoV-2 genome. Yousefi and co-workers (2020) [44] indicated that SARS-CoV-2 engaged a host of human miRNA including miR-424-5p that formed part of the crucial transforming growth factor beta (TGFB) signalling pathway. This pathway is associated with several cellular processes that include cell growth & differentiation, programmed cell death and cellular homeostasis. Demirci and Adan (2020) [45] reported that miR-6838-5p is one of the 21 different miRNAs with binding site within the SARS-coV-2 N (Nucleocapsid) protein encoded by this 1260 bp gene and is a contender for novel therapeutic solution.

The human microRNA miR-20b-3p [46] has been reported to overexpress in stromal cells and cure conditions resulting from oxalate deposition in mammals. Kidney is a common target for SARS-CoV-2. [47] In the kidney, ACE2 is expressed in the proximal tubules and somewhat with less intensity, in the glomeruli.[48] This provides logical evidence towards the kidney being one of the targets for SARS-CoV-2. Lesser reports of organ failure involving kidney may be in part due to the entry-point of the virus that makes kidney a distal destination along with the microRNA population that mounts protective action on the pathogen upon entry. This miRNA has two distinct seed sequences that encompass amino acid positions 1116 (Figure 3) & 1118 (Figure 4) respectively, of the S gene. Our bioinformatic analysis revealed two mutations, vizT1116T and D1118H altered the seed location of miR-20b-3p from this region (Figure 4). The former was reported from Austria while the latter, from Austria, USA and Italy respectively. Importantly, one of these mutations (T1116T) is synonymous in nature indicating a phenomenon similar to the mutation within the stop codon (position 1274) encountered by us for miR-511-3p. Here, the amino acid does not alter but a nucleotide does, resulting in abolition of seed sequence of a human miRNA. Similar alteration was also seen in a sequence submitted from Iran where a tactical synonymous mutation retained amino acid serine at position 659 but altered the seed sequence for miR-219a-1-3p.

In the canonical biogenesis pathway, the RNA Polymerase II transcribes miRNA primary precursors. This gradually leads to formation of mature miRNA that functions as a sequence-specific guide to trigger the effector complex that is known to bind to the 3’end of a gene to ensure regulation of translation and stability. [49] However, the concept of miRNA functioning by essentially binding to the 3’ untranslated region of a gene is an older dogma. Recent reports suggest that these non-coding RNA anneals within the ORF frames also mount desired gene regulation. [50] This is also substantiated by our study where over 11 miRNA lead sequences were identified within the open reading frame of the S glycoprotein gene of SARS-CoV-2.

The A845S mutation encountered by us in this study warrants a special mention. It was found to be the target of 7 different human miRNA. Guruprasad in an article published in January 2021 [51] analysed mutations prevalent in the spike protein of SARS-CoV-2. In this study 10333 spike protein sequences were studied and out of them, 8155 were found to harbour 1 or more mutations. In all, a total of 9654 mutations were encountered by the author that corresponded to 400 distinct mutation sites. However, A845S was not reported in this publication. A845S was reported to emerge in the second wave of COVID-19 pandemic in the USA.[52] It is tempting to conclude that perhaps an armour of miRNA along with other conditions encountered this virus since its emergence but subsequently failed with time during the late 2020 and early 2021 when this (and similar other) mutation emerged rendering over 7 different antiviral miRNA ineffective.

The emergence of synonymous mutations strategically erasing potential miRNA seed complementarity as observed in our study is intriguing. We detected a stop codon remaining unaltered by modifying a potent miRNA recognition site (Figure 2). We also detected similar synonymous mutations at position 845 and 1116 where the amino acids remain unaltered by the change affected potential binding site of a human miRNA (Figure 3 and Figure 4). Another interesting observation was alteration in miRNA seed sequence that is a target to multiple (n=7) different miRNA (Figure 1). It indicated a minimal genetic change by the pathogen in its genome by way of affecting a single base change that apparently brought in a magnified effect by deflecting several (n=7) miRNA, thereby preventing them from deregulating or destroying it.

Approaches are being adopted for designing a concoction of miRNA that can collectively target multiple regions of the SARS-CoV-2 not only at the 3’untranslated region but also within the open reading frame of the genes. [53] The delivery system can be non-viral in origin such as liposomes or other polymer-based carriers which are more suitable than viral vectors. This study throws important light on the viral adaptation by way of careful alteration of specific miRNA seed binding sites on its genes and pointing at the fact that an approach of a set of several miRNAs binding at multiple sites is only to be adopted by carefully avoiding potential mutational hotspots on the viral genome. Even then, gradual redundancy of these synthetic therapeutic small biomolecules might creep in due to selected mutation within miRNA seed locations within the viral genome as is demonstrated in our study using the S gene as a model.

## Reference

[1] Zhu N, Zhang D, Wang W, Li X, Yang B, Song J, Zhao X, Huang B, Shi W, Lu R, Niu P, Zhan F, Ma X, Wang D, Xu W, Wu G, Gao GF, Tan W; China Novel Coronavirus Investigating and Research Team. A Novel Coronavirus from Patients with Pneumonia in China, 2019. N Engl J Med. 2020 Feb 20;382(8):727–733.

[2] Rabi FA, Al Zoubi MS, Kasasbeh GA, Salameh DM, Al-Nasser AD. SARS-CoV-2 and Coronavirus Disease 2019: What We Know So Far. Pathogens. 2020 Mar 20;9(3):231

[3] Weiss SR, Leibowitz JL. Coronavirus pathogenesis. Adv Virus Res. 2011;81:85–164.

[4] Tang, Q., Song, Y., Shi, M., Cheng, Y., Zhang, W., & Xia, X. Q. Inferring the hosts of coronavirus using dual statistical models based on nucleotide composition. Scientific reports 5, 2015;17155.

[5] Wu C, Liu Y, Yang Y, Zhang P, Zhong W, Wang Y, Wang Q, Xu Y, Li M, Li X, Zheng M, Chen L, Li H. Analysis of therapeutic targets for SARS-CoV-2 and discovery of potential drugs by computational methods. Acta Pharm Sin B. 2020 May;10(5):766–788.

[6] Liu C, Zhou Q, Li Y, Garner LV, Watkins SP, Carter LJ, Smoot J, Gregg AC, Daniels AD, Jervey S, Albaiu D. Research and Development on Therapeutic Agents and Vaccines for COVID-19 and Related Human Coronavirus Diseases. ACS Cent Sci. 2020 Mar 25;6(3):315–331.

[7] Sanders JM, Monogue ML, Jodlowski TZ, Cutrell JB. Pharmacologic Treatments for Coronavirus Disease 2019 (COVID-19): A Review. JAMA. 2020 May 12;323(18):1824–1836.

[8] Lu R, Zhao X, Li J, Niu P, Yang B, Wu H, Wang W, Song H, Huang B, Zhu N, Bi Y, Ma X, Zhan F, Wang L, Hu T, Zhou H, Hu Z, Zhou W, Zhao L, Chen J, Meng Y, Wang J, Lin Y, Yuan J, Xie Z, Ma J, Liu WJ, Wang D, Xu W, Holmes EC, Gao GF, Wu G, Chen W, Shi W, Tan W. Genomic characterisation and epidemiology of 2019 novel coronavirus: implications for virus origins and receptor binding. Lancet. 2020 Feb 22;395(10224):565–574.

[9] Chen L, Liu W, Zhang Q, Xu K, Ye G, Wu W, Sun Z, Liu F, Wu K, Zhong B, Mei Y, Zhang W, Chen Y, Li Y, Shi M, Lan K, Liu Y. RNA based mNGS approach identifies a novel human coronavirus from two individual pneumonia cases in 2019 Wuhan outbreak. Emerg Microbes Infect. 2020 Feb 5;9(1):313–319.

[10] Chan JF, Kok KH, Zhu Z, Chu H, To KK, Yuan S, Yuen KY. Genomic characterization of the 2019 novel human-pathogenic coronavirus isolated from a patient with atypical pneumonia after visiting Wuhan. Emerg Microbes Infect. 2020 Jan 28;9(1):221–236.

[11] Letko M, Marzi A, Munster V. Functional assessment of cell entry and receptor usage for SARS-CoV-2 and other lineage B betacoronaviruses. Nat Microbiol. 2020 Apr;5(4):562–569.

[12] Fehr AR, Perlman S. Coronaviruses: an overview of their replication and pathogenesis. Methods Mol Biol. 2015;1282:1–23.

[13] Morse JS, Lalonde T, Xu S, Liu WR. Learning from the Past: Possible Urgent Prevention and Treatment Options for Severe Acute Respiratory Infections Caused by 2019-nCoV. Chembiochem. 2020 Mar 2;21(5):730–738.

[14] Trobaugh DW, Gardner CL, Sun C, Haddow AD, Wang E, Chapnik E, Mildner A, Weaver SC, Ryman KD, Klimstra WB. RNA viruses can hijack vertebrate microRNAs to suppress innate immunity. Nature. 2014 Feb 13;506(7487):245–248.

[15] Trobaugh DW, Klimstra WB. MicroRNA Regulation of RNA Virus Replication and Pathogenesis. Trends Mol Med. 2017 Jan;23(1):80–93.

[16] Scheel TK, Luna JM, Liniger M, Nishiuchi E, Rozen-Gagnon K, Shlomai A, Auray G, Gerber M, Fak J, Keller I, Bruggmann R, Darnell RB, Ruggli N, Rice CM. A Broad RNA Virus Survey Reveals Both miRNA Dependence and Functional Sequestration. Cell Host Microbe. 2016 Mar 9;19(3):409–23.

[17] Trobaugh DW, Klimstra WB. MicroRNA Regulation of RNA Virus Replication and Pathogenesis. Trends Mol Med. 2017 Jan;23(1):80–93.

[18] Barbu MG, Condrat CE, Thompson DC, Bugnar OL, Cretoiu D, Toader OD, Suciu N, Voinea SC. MicroRNA Involvement in Signaling Pathways During Viral Infection. Front Cell Dev Biol. 2020 Mar 10;8:143.

[19] Bernier A, Sagan SM. The Diverse Roles of microRNAs at the Host-Virus Interface. Viruses. 2018 Aug 19;10(8):440.

[20] Zhang H, Rostami MR, Leopold PL, Mezey JG, O’Beirne SL, Strulovici-Barel Y, Crystal RG. Expression of the SARS-CoV-2 ACE2 Receptor in the Human Airway Epithelium. Am J Respir Crit Care Med. 2020 Jul 15;202(2):219–229.

[21] Sievers, Fabian; Barton, Geoffrey J.; Higgins, Desmond G., Multiple Sequence Alignments. Bioinformatics. editor / Andreas D. Baxevanis; Gary D. Bader; David S. Wishart. 4. ed. Wiley, 2020 pp. 227–250.

[22] UniProt Consortium. UniProt: the universal protein knowledgebase in 2021. Nucleic Acids Res. 2021 Jan 8;49(D1):D480–D489.

[23] Yuhao Chen, Xiaowei Wang, miRDB: an online database for prediction of functional microRNA targets, Nucleic Acids Research, 08 January 2020;48(D1): D127–D131.

[24] Rampal, L. & Boon Seng, L. Coronavirus disease (COVID-19) pandemic. Med J Malaysia, March 2020;Vol. 75, No 2:95–97.

[25] Kandeel M, Ibrahim A, Fayez M, Al-Nazawi M. From SARS and MERS CoVs to SARS-CoV-2: Moving toward more biased codon usage in viral structural and nonstructural genes. J Med Virol. 2020 Jun;92(6):660–666.

[26] Baig AM, Khaleeq A, Ali U, Syeda H. Evidence of the COVID-19 Virus Targeting the CNS: Tissue Distribution, Host-Virus Interaction, and Proposed Neurotropic Mechanisms. ACS Chem Neurosci. 2020 Apr 1;11(7):995–998.

[27] Sv P, Lathabhavan R, Ittamalla R. What concerns Indian general public on second wave of COVID-19? A report on social media opinions. Diabetes Metab Syndr. 2021 Apr 14;15(3):829–830.

[28] Ylösmäki E, Hakkarainen T, Hemminki A, Visakorpi T, Andino R, Saksela K. Generation of a conditionally replicating adenovirus based on targeted destruction of E1A mRNA by a cell type-specific MicroRNA. Journal of Virology. 2008;82(22):11009–11015.

[29] Barnes D, Kunitomi M, Vignuzzi M, Saksela K, Andino R. Harnessing endogenous miRNAs to control virus tissue tropism as a strategy for developing attenuated virus vaccines. Cell Host Microbe. 2008 Sep 11;4(3):239–48.

[30] Otsuka M, Jing Q, Georgel P, New L, Chen J, Mols J, Kang YJ, Jiang Z, Du X, Cook R, Das SC, Pattnaik AK, Beutler B, Han J. Hypersusceptibility to vesicular stomatitis virus infection in Dicer1-deficient mice is due to impaired miR24 and miR93 expression. Immunity. 2007 Jul;27(1):123–34.

[31] Davis M, Sagan SM, Pezacki JP, Evans DJ, Simmonds P. Bioinformatic and physical characterizations of genome-scale ordered RNA structure in mammalian RNA viruses. J Virol. 2008 Dec;82(23):11824–36.

[32] Simmons G, Gosalia DN, Rennekamp AJ, Reeves JD, Diamond SL, Bates P. Inhibitors of cathepsin L prevent severe acute respiratory syndrome coronavirus entry. Proc Natl Acad Sci U S A. 2005 Aug 16;102(33):11876–81.

[33] Glowacka I, Bertram S, Müller MA, Allen P, Soilleux E, Pfefferle S, Steffen I, Tsegaye TS, He Y, Gnirss K, Niemeyer D, Schneider H, Drosten C, Pöhlmann S. Evidence that TMPRSS2 activates the severe acute respiratory syndrome coronavirus spike protein for membrane fusion and reduces viral control by the humoral immune response. J Virol. 2011 May;85(9):4122–34.

[34] Matsuyama S, Nagata N, Shirato K, Kawase M, Takeda M, Taguchi F. Efficient activation of the severe acute respiratory syndrome coronavirus spike protein by the transmembrane protease TMPRSS2. J Virol. 2010 Dec;84(24):12658–64.

[35] Shulla A, Heald-Sargent T, Subramanya G, Zhao J, Perlman S, Gallagher T. A transmembrane serine protease is linked to the severe acute respiratory syndrome coronavirus receptor and activates virus entry. J Virol. 2011 Jan;85(2):873–82.

[36] Kawase M, Shirato K, van der Hoek L, Taguchi F, Matsuyama S. Simultaneous treatment of human bronchial epithelial cells with serine and cysteine protease inhibitors prevents severe acute respiratory syndrome coronavirus entry. J Virol. 2012 Jun;86(12):6537–45.

[37] Finnerty JR, Wang WX, Hébert SS, Wilfred BR, Mao G, Nelson PT. The miR-15/107 group of microRNA genes: evolutionary biology, cellular functions, and roles in human diseases. J Mol Biol. 2010 Sep 24;402(3):491–509.

[38] Peng S, Wang J, Wei S, Li C, Zhou K, Hu J, Ye X, Yan J, Liu W, Gao GF, Fang M, Meng S. Endogenous Cellular MicroRNAs Mediate Antiviral Defense against Influenza A Virus. Mol Ther Nucleic Acids. 2018 Mar 2;10:361–375.

[39] Morales L, Oliveros JC, Fernandez-Delgado R, tenOever BR, Enjuanes L, Sola I. SARS-CoV-Encoded Small RNAs Contribute to Infection-Associated Lung Pathology. Cell Host Microbe. 2017 Mar 8;21(3):344–355.

[40] Nersisyan, S., Engibaryan, N., Gorbonos, A., Kirdey, K., Makhonin, A., & Tonevitsky, A.. Potential role of cellular miRNAs in coronavirus-host interplay. PeerJ. 2020; 8, e9994.

[41] Zheng, C., Zheng, Z., Sun, J. et al. MiR-16-5p mediates a positive feedback loop in EV71-induced apoptosis and suppresses virus replication. Sci Rep 7. 2017, 16422 (2017).

[42] Kim WR, Park EG, Kang KW, Lee SM, Kim B, Kim HS. Expression Analyses of MicroRNAs in Hamster Lung Tissues Infected by SARS-CoV-2. Mol Cells. 2020 Nov 30;43(11):953–963.

[43] Jafarinejad-Farsangi S, Jazi MM, Rostamzadeh F, Hadizadeh M. High affinity of host human microRNAs to SARS-CoV-2 genome: An in silico analysis. Non-coding RNA Research. 2020 Dec;5(4):222–231

[44] Yousefi H, Poursheikhani A, Bahmanpour Z, et al. SARS-CoV infection crosstalk with human host cell noncoding-RNA machinery: An in-silico approach. Biomedicine & Pharmacotherapy = Biomedecine & Pharmacotherapie. 2020 Oct;130:110548.

[45] Saçar Demirci MD, Adan A. Computational analysis of microRNA-mediated interactions in SARS-CoV-2 infection. PeerJ. 2020 Jun 5;8:e9369.

[46] Shi J, Duan J, Gong H, Pang Y, Wang L, Yan Y. Exosomes from miR-20b-3p-overexpressing stromal cells ameliorate calcium oxalate deposition in rat kidney. J Cell Mol Med. 2019 Nov;23(11):7268–7278.

[47] Hardenberg JB, Luft FC. Covid-19, ACE2 and the kidney. Acta Physiol (Oxf). 2020 Sep;230(1):e13539.

[48] Mizuiri S, Ohashi Y. ACE and ACE2 in kidney disease. World J Nephrol. 2015 Feb 6;4(1):74–82.

[49] Girardi E, López P, Pfeffer S. On the Importance of Host MicroRNAs During Viral Infection. Front Genet. 2018 Oct 2;9:439.

[50] Li H. Exploration of Alternative Mechanism for MiR-596-mediated Down-regulation of LGALS3BP in Oral Squamous Cell Carcinoma. Kokubyo Gakkai zasshi. The Journal of the Stomatological Society, Japan. 2015 Jul;82(2):55–61.

[51] Guruprasad L. Human SARS CoV-2 spike protein mutations. Proteins. 2021 May;89(5):569–576.

[52] Long, S. W., Olsen, R. J., Christensen, P. A., Bernard, D. W., Davis, J. J., Shukla, M., Nguyen, M., Saavedra, M. O., Yerramilli, P., Pruitt, L., Subedi, S., Kuo, H. C., Hendrickson, H., Eskandari, G., Nguyen, H., Long, J. H., Kumaraswami, M., Goike, J., Boutz, D., Gollihar, J., … Musser, J. M.. Molecular Architecture of Early Dissemination and Massive Second Wave of the SARS-CoV-2 Virus in a Major Metropolitan Area. mBio, 11(6).2020; e02707–20:1–30.

[53] El-Nabi SH, Elhiti M, El-Sheekh M. A new approach for COVID-19 treatment by microRNA. Med Hypotheses. 2020 Oct;143:110203.

[54] Lewis BP, Burge CB, Bartel DP. Conserved seed pairing, often flanked by adenosines, indicates that thousands of human genes are microRNA targets. Cell. 2005 Jan 14;120(1):15–20.

